# Predicting metastasis with a novel biophysical cell-adhesion force technique

**DOI:** 10.1101/2020.04.13.026526

**Authors:** Jessie Gan, Zhao Zhihai, Yu Miao

## Abstract

Metastasis is widely accepted to be responsible for approximately 90% of all cancer deaths. Current research on metastasis prediction often centers on gene sequencing; however, these analyses must account for the complexity of gene regulation and rely on comprehensive datasets. To investigate the process from a simpler, non-genomic angle, some studies indicate differences in cell adhesion force, an important physical process in metastasizing cells. However, cell adhesion force methods tend to focus on cell population approaches and therefore have their drawbacks in cost or efficiency, rendering them impractical outside a research setting. In this work, we test a novel and inexpensive bead-pipette assay to investigate the adhesion forces of non-metastatic NIH3T3 cells and mutated RasV12 cells, a metastatic model cell line.

Control cells and RasV12 cells were evaluated with wound healing, spreading area, and focal adhesion (FA) analysis assays. Then cells were tested by the novel bead-pipette assay, which uses a fibronectin-coated bead and a glass micropipette to measure cell adhesion force using Hooke’s law.

The RasV12 cells had faster migration, polarized cell shape, and smaller FA area than control cells. The RasV12 cells also exerted higher adhesion forces than control cells and a potential force threshold was determined for distinguishing metastatic cells through a Receiver Operating Characteristic (ROC) curve. An ROC curve was computed for all other assays and the bead-pipette assay was shown to perform higher as a classifier than other assays.

The RasV12 cells had increased metastatic potential compared to control. The novel bead-pipette assay showed potential as a classifier for determining metastasizing cells from non-metastatic cells. With further work, it may serve as a clinical diagnostic tool for cancer patients or as a testbed to be used in the development of anti-metastatic drugs.

## Introduction

Metastasis, the migration of cancer cells to a secondary tumor location, is a significant contributor to cancer patient deaths(1). The onset and progression of metastasis is difficult to predict and as yet, no universal prognostic metastasis marker has been identified. Most research is focused on genomic markers through sequencing or microarray assays, but results are not comprehensive and are typically cancer-type specific(2–5). Diagnostics using sequencing data paired with machine-learning models, although getting faster and cheaper, still must account for the complexity of the gene regulation of metastasis due to factors such as alternative splicing, post-translational modifications, and protein processing(6,7). In addition, these diagnostic models must be trained on immense and comprehensive datasets(8–10), which are tedious to curate. Other tests include blood marker testing, CT scans, and MRI, which cannot diagnose metastasis until the tumor has already metastasized(11).

Current literature indicates that cell adhesion force plays a major role in metastasis and is influenced by cell genotype(12–15). Metastasis is defined by invasion and motility of the cancer cell from the home base to a secondary location. Cell motility involves the integration of multiple mechanical and chemical cues, many of which are driven by the adhesion of the cell to the local extracellular matrix in its immediate neighborhood(16,17). Given that metastasizing cells are known to move actively through the extracellular matrix and are affected by environmental forces(18–20), it is possible that their adhesion forces will differ from those of stationary cells. In the interest of simplifying and accelerating the detection of metastatic cancers, using adhesion force as a metric to differentiate non-metastatic and metastatic cells could be a low-cost and high-throughput alternative to diagnostics based on genetic sequencing.

Metastasis is involved with cell adherence to extracellular matrix (ECM) and other cells through protein complexes called focal adhesions (FAs)(21–23). FAs, the primary focus of this work, are the interface for cells to interact and sense their local microenvironment, and they are central hubs for mechanotransduction, ECM sensing, and directing cell migration(23–26). Sites of FAs are initiated by the binding of integrin receptor proteins to ECM components and the subsequent recruitment and clustering of cytoplasmic proteins and cytoskeletal elements. Integrin, a transmembrane protein in FAs, has been shown to have a significant role in metastatic processes(22,27,28), and it binds to collagen and fibronectin (ECM components) through hydrogen bonds and metal coordination(29–31). In stationary cells, the initial, nascent FAs mature into larger, established FAs, which provide passive anchorage(32). However, in motile cells, the cytoplasmic components generate a pulse of traction force upon the ECM substrate, such as collagen or fibronectin (FN), then disassemble to form new FAs to propel the cell forward(33).

Motile cells are often observed to be polarized and have distinct leading and trailing edges(36,37). The leading edge, characterized by the direction of movement, is formed by protrusions controlled through actin polymerization. The leading edge is also characterized by the FA turnover rate, the rate with which FAs assemble and disassemble to form new FAs(23,38). In motile cells, the leading edge may have a high turnover rate of FAs to continue adhering to ECM as the cell moves forward(38).

The idealized workflow is where patient biopsy samples would be directly tested in the hospital lab to test patients’ cells for metastasis, and thereby providing a valuable insight in the progression of the disease. The adhesion of the cells obtained from the biopsy adhesion would be tested by an adhesion force technique and classified based on a force threshold. In addition, drugs tested for metastatic prevention can be evaluated for their relative effectiveness with the adhesion force on certain extracellular substrates.

In order to use cell adhesion as a metric, there must be a consistent, versatile, and affordable technique for measuring it. Many methods have been developed for quantifying cell adhesion, such as traction force microscopy(39), centrifugal force assays(40), atomic force microscopy(41), and single-cell aspiration(42). However, although many have advantages including specific force observation and standard reproducibility, they also have disadvantages such as low maximum forces and inaccurate modeling due to cell or chamber deformation(42,43). They also can be expensive or require an extensive number of cells, rendering them inviable in a clinical setting where they could be assisting the diagnosis of cancer patients.

To address the limitations of current methods for measuring cell adhesion, the Yan Jie lab has developed the bead-pipette assay, a single-cell manipulation method of measuring adhesion force implemented in this work. Its advantages include inexpensive materials, efficient measurements, and precise control, and it shows potential to be not only applicable in a research setting but also in a clinical and translational environment.

Currently metastasis accounts for an overwhelming majority of cancer deaths worldwide(1), and an integral component in metastasizing cells is their adhesion strength based on the establishment, maturation rate, and deconstruction of FAs. In this work, we study the potential to use cell adhesion as a metric for differentiating between metastatic and non-metastatic cancers with a novel, biophysical assay. This is first investigated by first studying the molecular interactions between integrins and FN, and the role of FAs in cell adhesion. Next we developed an experiment to test the viability of the bead-pipette assay as a technique to measure differentiating cell adhesion force, and we also assess other phenotypic aspects of the cells, such as migration distance and FA size, to investigate their relationship to cell motility.

We hypothesized that the bead-pipette can effectively differentiate between metastatic and non-metastatic cells, and that this approach is suitable for identifying metastasizing cells in patient samples through cell adhesion force. The implementation of this technique in a clinical setting could present a simple solution to the diagnosis of metastasis, by applying a physics solution to a biological problem in an interdisciplinary application.

## Methods

In this work, we use p53-knockout, mouse fibroblast NIH3T3 cells from the American Type Culture Collection as control, non-metastatic cells. we use RasV12-transformed NIH3T3 cells as metastatic cells(44). The RasV12 cancer model activates metastatic-related pathways that promote processes such as cell proliferation and invasion(44–46), making it suitable for this work. The mutation results in a perpetually active Ras-GTP complex that is unable to be inactivated by the Ras-GTPase activating proteins (GAPs)(47), and thereby continues to upregulate metastatic-related pathways.

### Wound Healing Assay

We performed the wound healing assay to gauge the initial metastatic potential of the cells. we first seeded the cells on collagen-coated glass with dividers from Ibidi GmbH (Germany) for 24 hours until 100% confluency was reached. Then we removed the dividers and imaged the cells for another 24 hours afterwards as they moved to cover the gap from the divider. We analyzed data at the eight-hour mark in the corresponding videos at 10x magnification under a light microscope, and calculated the distance migrated in micrometers through the Fiji ImageJ visualization program(48).

### Spreading Area

We calculated the spreading area of the cells through imaging the cells with a light microscope at 20x magnification. We photographed the cells at a low confluence and quantified their area by outlining the cell membranes in Fiji ImageJ visualization program(48). We performed this at n=20 for each condition, for a total of 40 measurements.

### Focal Adhesions

We measured the focal adhesion area through immunofluorescence visualization. Cells grown over two weeks were seeded overnight on small petri-dishes with a FN-coated glass well. We used a 4% paraformaldehyde solution to fix the cells, then 0.2% Triton to perforate the cell membrane. We added Bovine Serum Albumin, a blocker to prevent non-specific binding, to the fixed and permeabilized cells. We then added a primary antibody for paxillin from Cell Signaling Technology and incubated the cells at 4 °C overnight. The next day we added a secondary antibody conjugated with Green Fluorescence Protein for the primary antibody. We stained the nuclei with 4’,6-diamidino-2-phenylindole (DAPI). All reagents were purchased from Thermo Fisher Scientific (Waltham, MA) unless otherwise noted.

We performed fluorescence microscopy was performed with a Nikon A1R confocal microscope. We visualized the cells under a 405 and 488 nm wavelength laser for the nuclei and anti-paxillin antibody, respectively. We identified focal adhesions were identified through setting a brightness threshold of 3320 gray level in a 16 bit image (gray level range of 0-65535), and then quantified them by the “Analyze Particles” feature in Fiji ImageJ(48). FA measurements were taken for n=20 for each condition, for a total of 40 cells.

### Force Quantification - Bead Pipette Assay

The force quantification bead assay is a novel force-measurement method that utilizes a ECM-coated bead and a glass micropipette to measure cell adhesion force. Cells were seeded overnight in a chamber composed of a polydimethylsiloxane cutout between two glass slides coated in collagen. Before measurement, FN-coated beads made in the lab were added to the chamber. The FN-coated beads were amino-coated polybeads, incubated with glutaraldehyde, and then with FN.

1 mm glass micropipettes were pulled to a fine point of about 2 µm in diameter, and were attached to a small water reservoir, through which suction into the pipette could be controlled by changing the relative height to the microscope stage. The pipette was maneuvered into the cell chamber with a micromanipulator, and was positioned to attach a bead by suction force. The cell chamber was visualized with an Olympus Live EZ microscope under a 20x air lens.

Using the micropipette to manipulate the FN-coated bead, cells were tested for adhesion force by bringing the bead in contact with the cell surface and allowing the integrin-FN interactions and FAs to form for 1 minute. Then the cell was moved in the horizontal direction at a speed of 10 μm/sec by moving the microscope stage until the bead broke contact with the cell. The process was photographed every 5 μm. After the bead broke contact with the cell surface, the rebound distance of the pipette was measured in Fiji ImageJ(48) and the force was calculated through the displacement and the spring constant of the micropipette, as shown in Figure 5A, B.

This procedure was performed at n=20 for each condition, for a total of 40 cells. The micropipette spring constant was calculated in the lab. The corresponding spring constant and distance were used to calculate the force values.

### Statistical Analysis

For each assay the Student’s T-Test(44) was calculated to determine if the difference between the control and RasV12 cells was statistically significant. The difference was considered statistically significant if the probability value (p-value) was below 0.05.

To test the performance of adhesion force as a binary classifier of metastatic and benign cells, the statistical analyses of an initial confusion matrix and a Receiver Operating Characteristic (ROC) curve were computed in scikit-learn(45). The confusion matrix is a grid that displays the percentage of true positives, true negatives, false positives, and false negatives the classifier produced based on an arbitrary force threshold, which was initially chosen as the lower standard deviation value of the RasV12 forces. To optimize an appropriate threshold and to compare the adhesion force as a classifier against other features measured, such as spreading area, focal adhesions, and cell migration, ROC curves of the rate of false positives vs. the rate of true positives were calculated. The area under the ROC curves and accuracy, precision, recall, and Cohen’s Kappa of the optimal threshold were calculated in Scikit-Learn(45).

## Results and Discussion

The wound healing assay revealed that the RasV12 cells migrate farther than non-mutated NIH3T3 cells over a period of 8 hours, and farther migration is a characteristic of metastatic cells (Fig 2A). However, the control cells tend to move together while the RasV12 cells move independently (Fig 2B). The control cells appear to have stronger cell-cell adhesion through proteins such as cadherin, whereas the RasV12 cells are separated and seem to have weaker cell-cell adhesion, which is another characteristic of metastatic cells.

**Fig 1.**
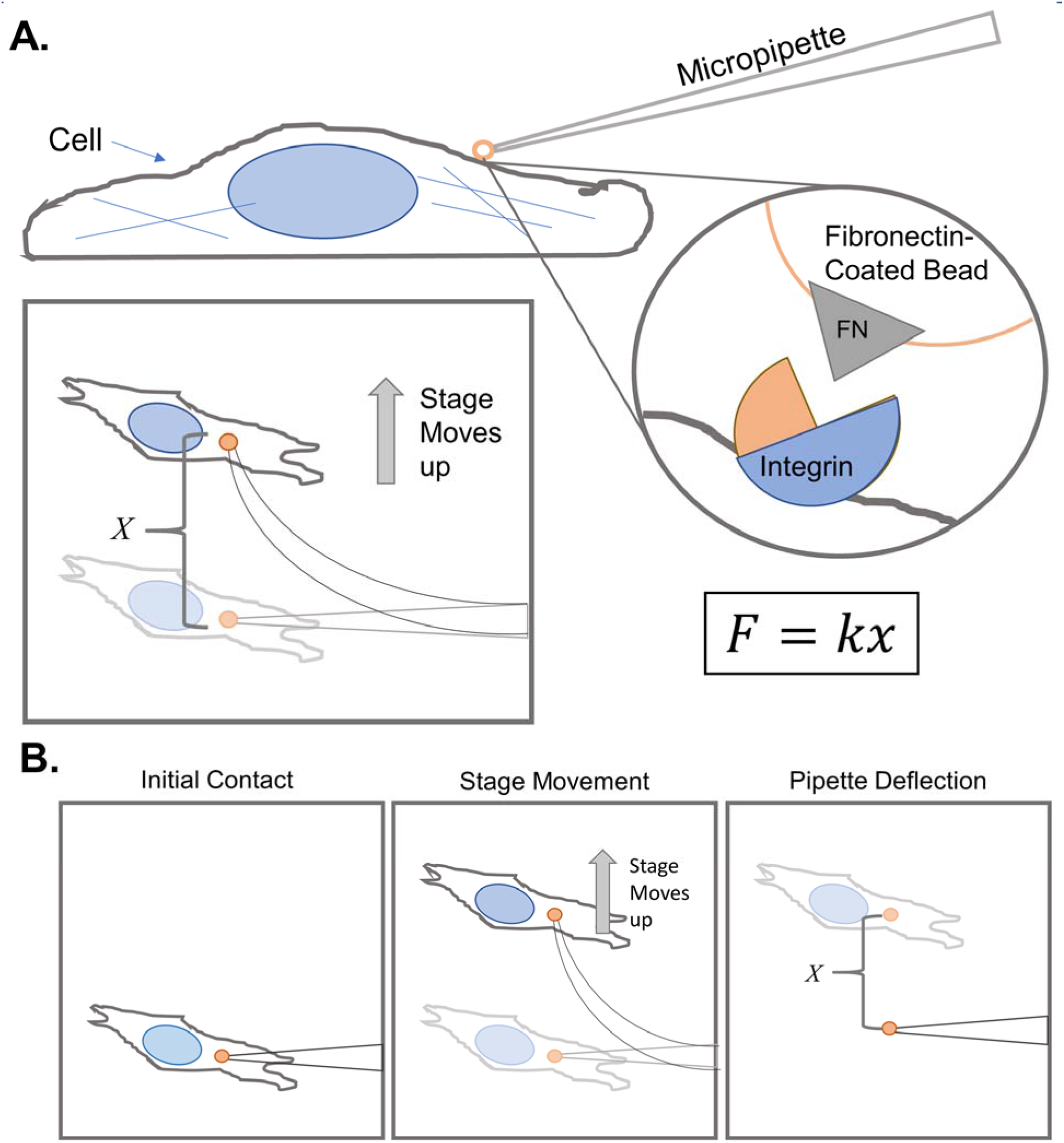
The bead-pipette method, using a flexible micropipette and FN-coated bead. **A.** The force is quantified by contacting the bead with the cell, then measuring the deflection of the micropipette when the cell is moved away at 10 μm/sec in increments of 5 μm. Multiplying the deflection distance (x) by the spring constant (k) of the pipette gives the force needed to break the integrin-FN interaction. **B.** A sequential schematic of the bead-pipette assay steps from an aerial view. First the bead is lowered onto the surface of the cell at Initial Contact, then the stage is incrementally raised in the horizontal direction by the microscope. When the bead breaks contact with the cell the distance is measured with image processing software Fiji ImageJ(43).

**Fig 2A.**
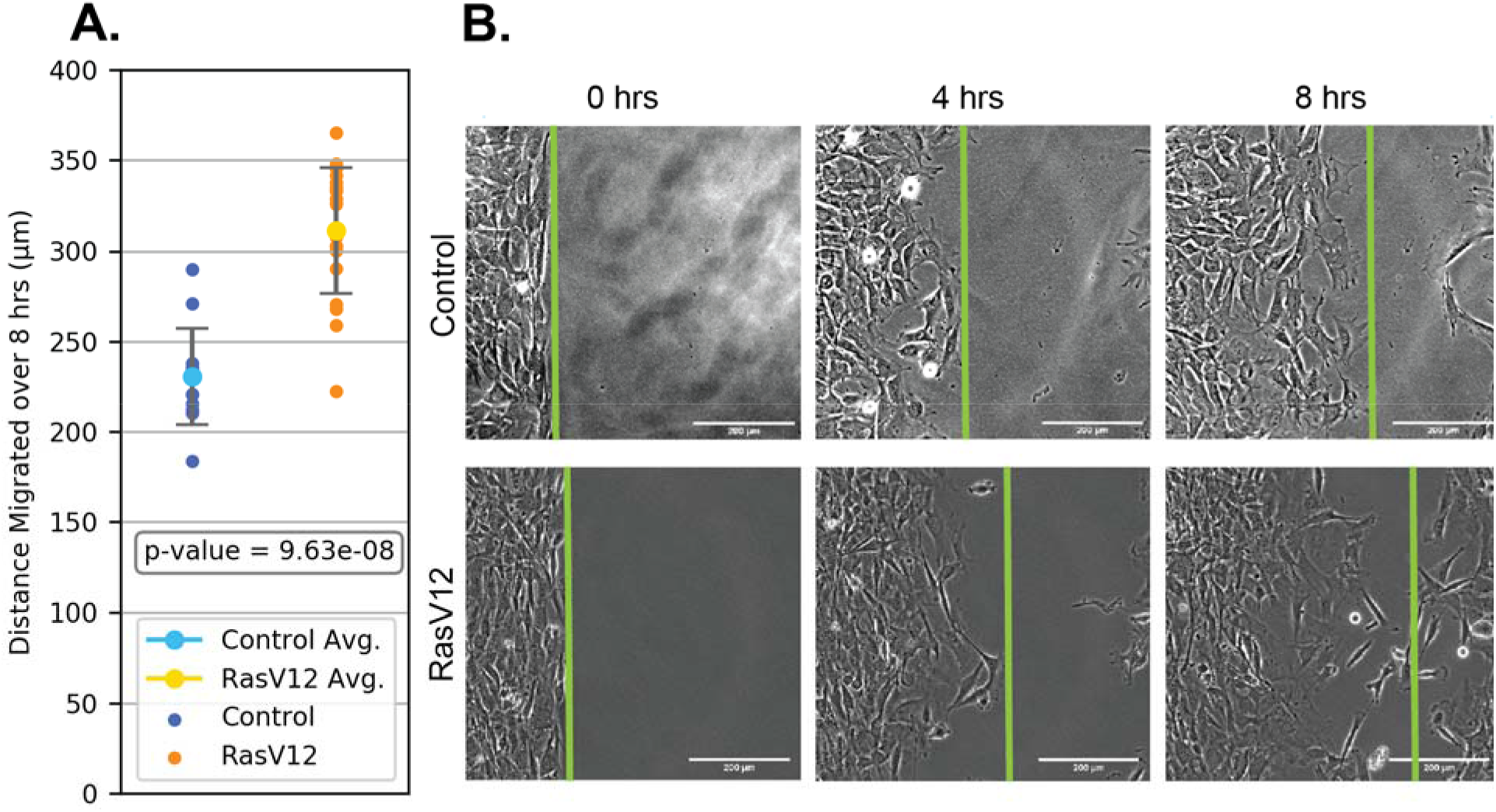
Wound healing assay shows that the RasV12 cells had a higher migration potential than the non-mutated NIH3T3 cells. Over a period of 8 hours the RasV12 cells migrated farther. The control cells averaged at 230.45 μm with ±26.76 error, and the RasV12 cells averaged at 311.35 μm with ±34.62 error. Plot made with python data visualization package Matplotlib(46). **B.** Photographs show that the RasV12 phenotype moves farther than the control phenotype.

The RasV12 phenotype cells have significantly less spreading area than the control cells in Fig 3A. Representative cells are shown in Fig 3B, where the control cells are larger, with tendrils of lamellipodium anchoring them over an extensive area. Their lack of polarization and large area do not indicate a specific direction. In contrast, the RasV12 cells are thinner and tapered, showing polarization of the FAs to a leading edge, a characteristic of migrating cells. The leading edge would likely have nascent FAs forming, and the cell would direct a backwards force, pulling itself along the extracellular matrix, perpetually forming new FAs at its leading edge and dissembling them at the trailing edge(27,30). The polarization of the RasV12 cells indicate their direction of movement, and result in a largely different cell shape than control cells.

**Fig 3A.**
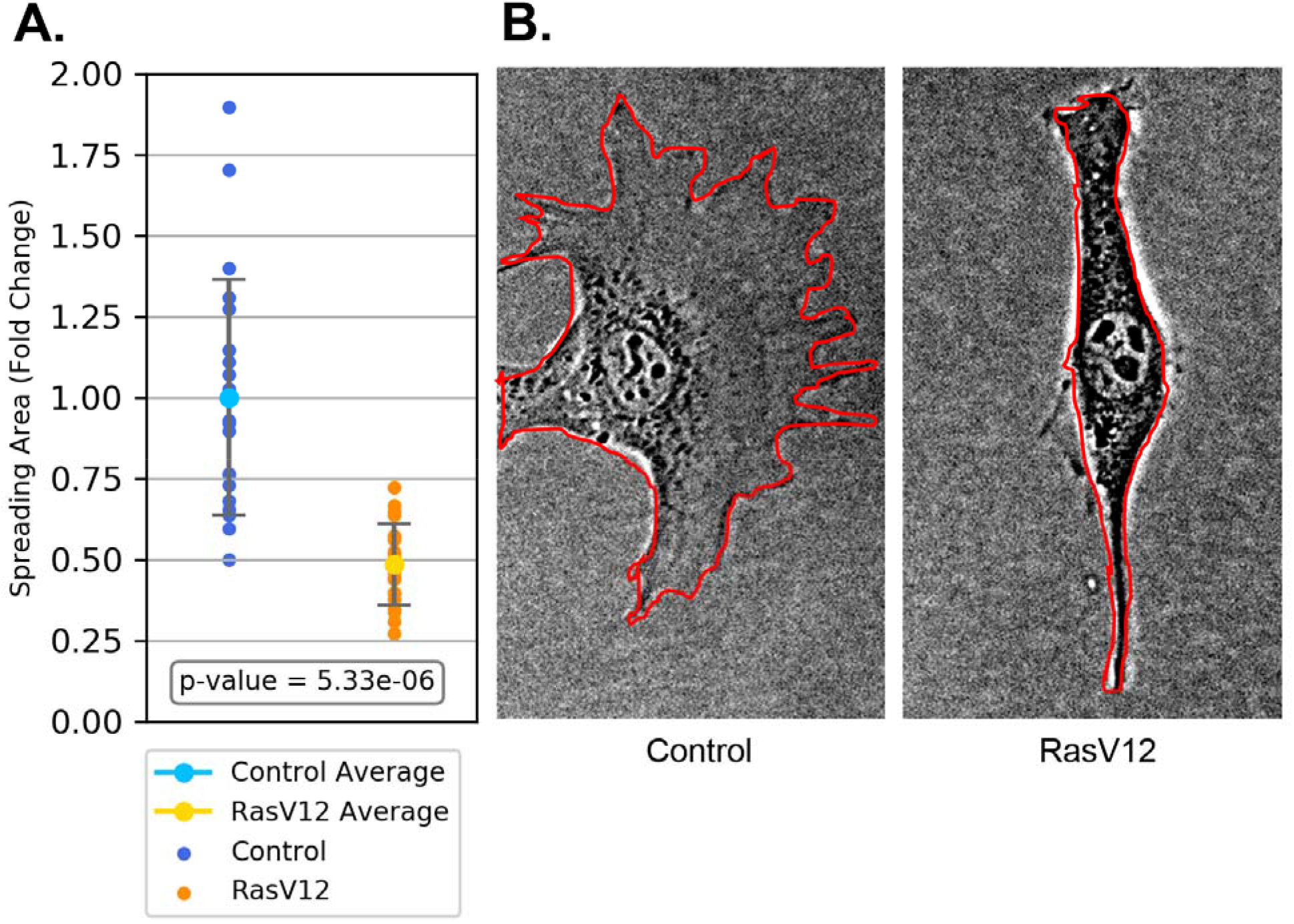
The RasV12 cells show less spreading area than the control cells. The data is normalized to the mean of the control cell area. The plot was made with Matplotlib(46). **B.** Spreading area outlines. These are representative examples of control and RasV12 cells with their areas outlined in Fiji ImageJ(43).

The RasV12 cells overall showed fewer FAs (Fig 4A), coupled with less total FA area (Fig 4B) and smaller individual FA size (Fig 4C). This indicates less anchorage to their substrate than control cells. Motile cells will likely require fewer and smaller, nascent FAs; a moving cell needs to swiftly synthesize and deconstruct FAs. These results imply the migratory potential of the RasV12 cells. On the other hand, the control cells have larger FAs and more FA area. These features are characteristic of mature FAs, which are more established instead of transient(27). More representative cells are shown in Fig 5.

**Fig 4.**
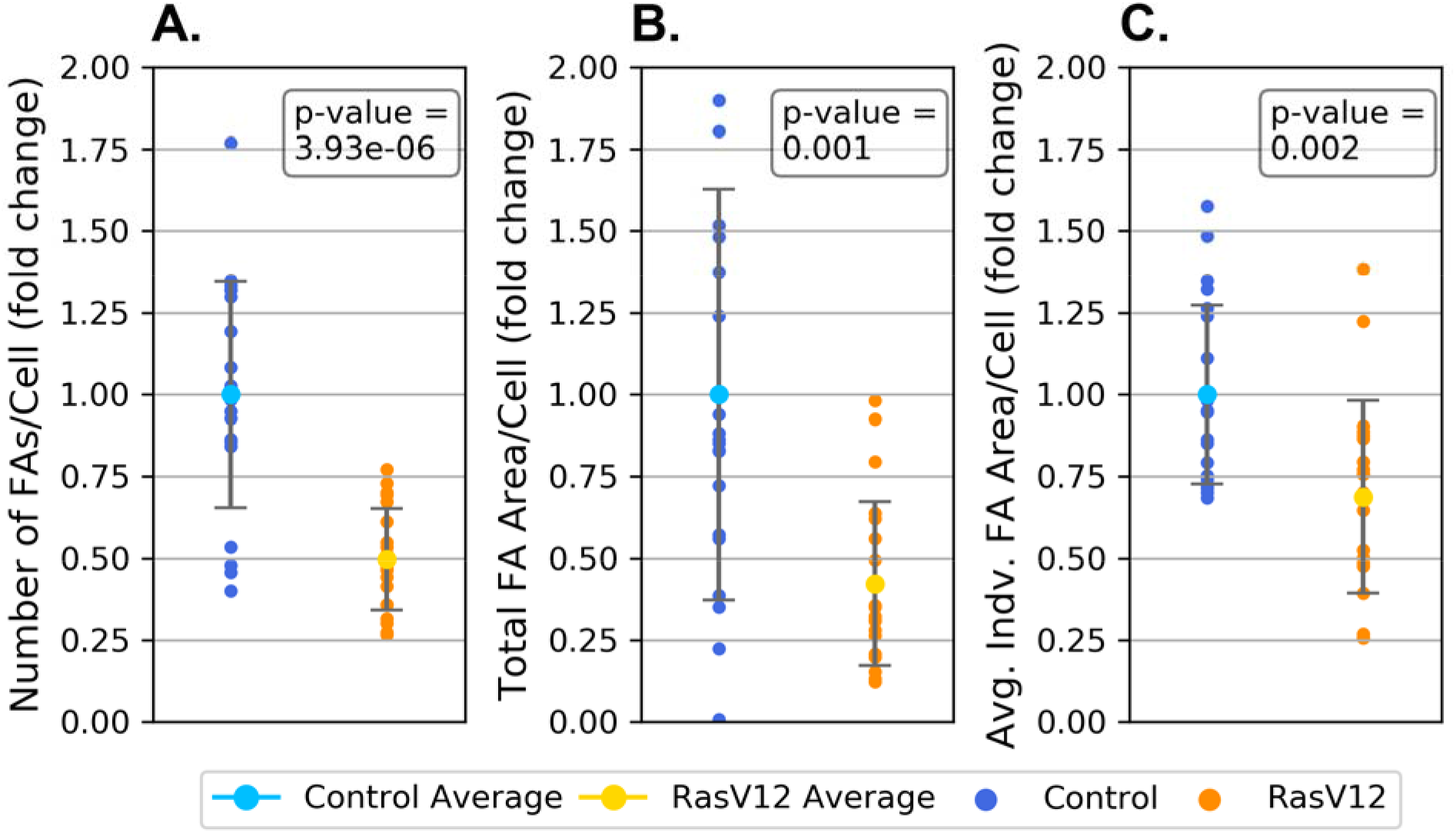
Focal adhesion quantitative analysis shows that RasV12 cells have less and smaller FAs than control. Control cells are in blue and RasV12 cells are in orange. All plots were made with Matplotlib(46) **A.** RasV12 cells have less total FAs than control cells. The data is normalized to the mean of the control number of FAs per cell. **B.** RasV12 cells have less total FA area than control cells. This data is normalized to the mean of the control cell focal adhesion size. **C.** RasV12 cells have smaller individual FA areas than control cells. This data is normalized to the average individual FA size for control cells.

**Fig 5.**
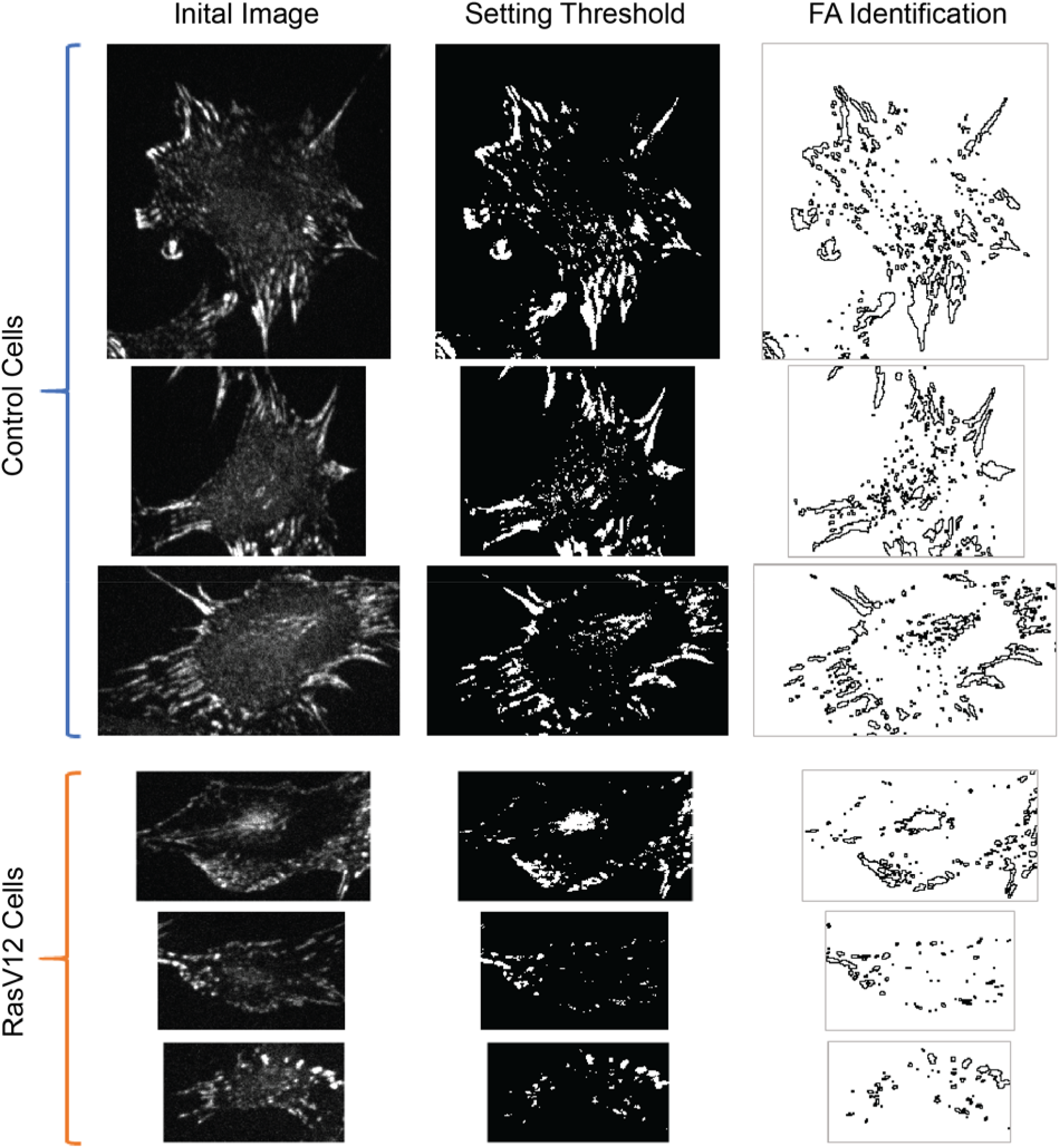
Representative cells are shown for each condition and each step in the process of calculating FAs through immunofluorescence assay in Fiji ImageJ(43). The original images were automatically adjusted to view and section off individual cells. Then a standard threshold was set and kept among all samples, and finally the “Analyze Particles” function was utilized to identify individual FAs.

In the bead-pipette assay, the RasV12 cells adhered to the FN-coated beads stronger than the control cells (Fig 6A). On average, they adhered about twice as strong. From the FA analysis, the RasV12 cells are shown to establish small FAs, and these nascent FAs appear to yield high adhesion force regardless of their size. On the other hand, the control cells are likely synthesizing FAs that form to maturity, however these are slowly assembled and deconstructed(23), and therefore may exert low forces when initially forming. The stages of the bead-pipette assay are shown in Fig 6B, where the bead is lowered onto the surface of the cell and a noticeably larger deflection is recorded for the RasV12 cells, indicating a higher force with similar micropipette spring constants.

**Fig 6.**
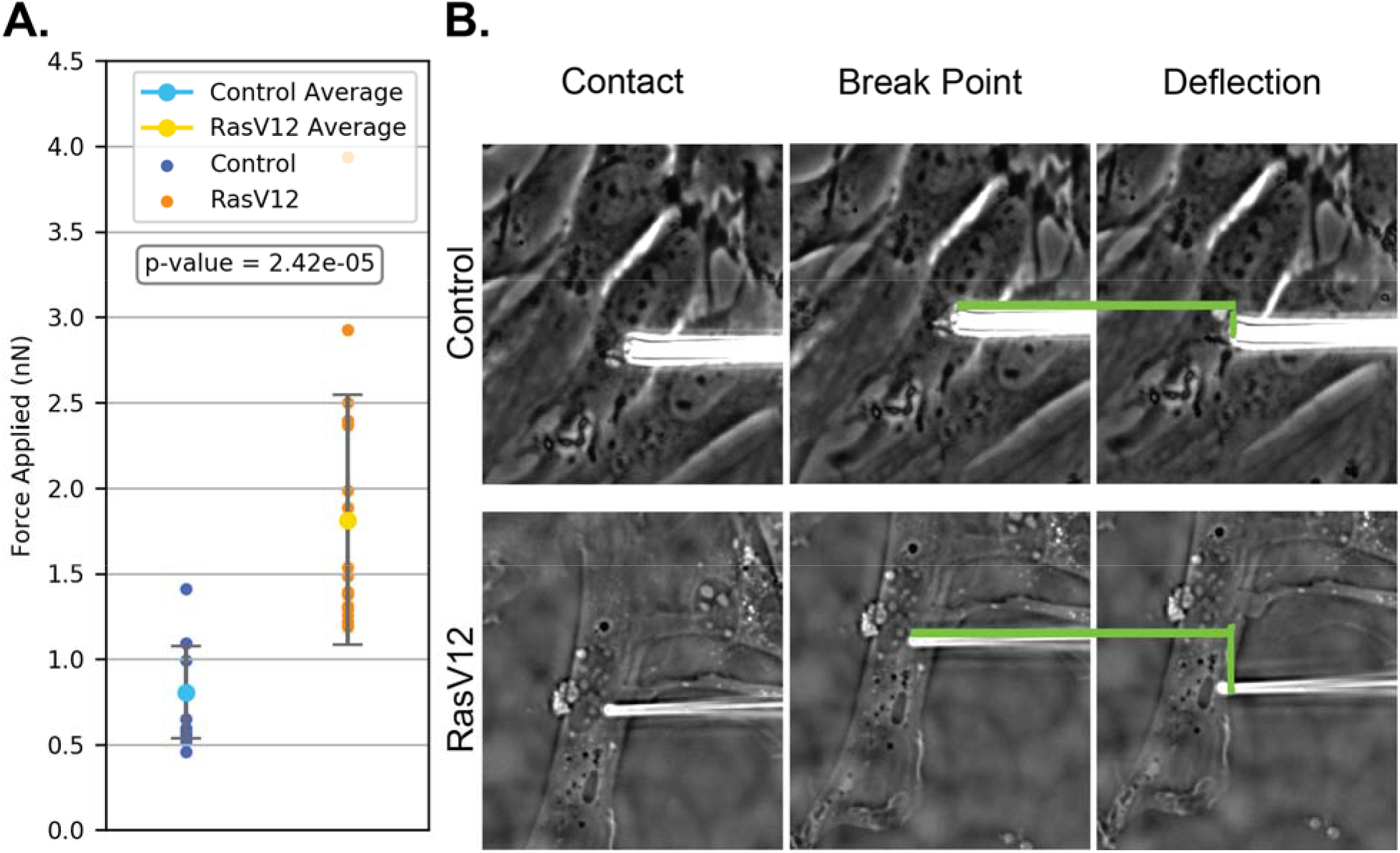
**A.** RasV12 Cells’ FAs exert higher force on FN ECM than control cells. The RasV12 phenotype cells have significantly higher force than the control cells. Plot was made in Matplotlib(46). **B.** Difference in pipette deflection between control and RasV12 cells in the bead assay. A larger distance contributes to a larger force exerted, in combination with the measured stiffness, or spring constant.

Statistical analysis through a confusion matrix was performed for the adhesion force classifier, where cells were considered metastasizing if their adhesion force was above the lower standard deviation value from average of the RasV12 forces (Fig 7A). This force threshold was able to account for all RasV12 cells, but some control cells were able to exert a force in the range of metastasizing cells, resulting in false positive values.

**Fig 7.**
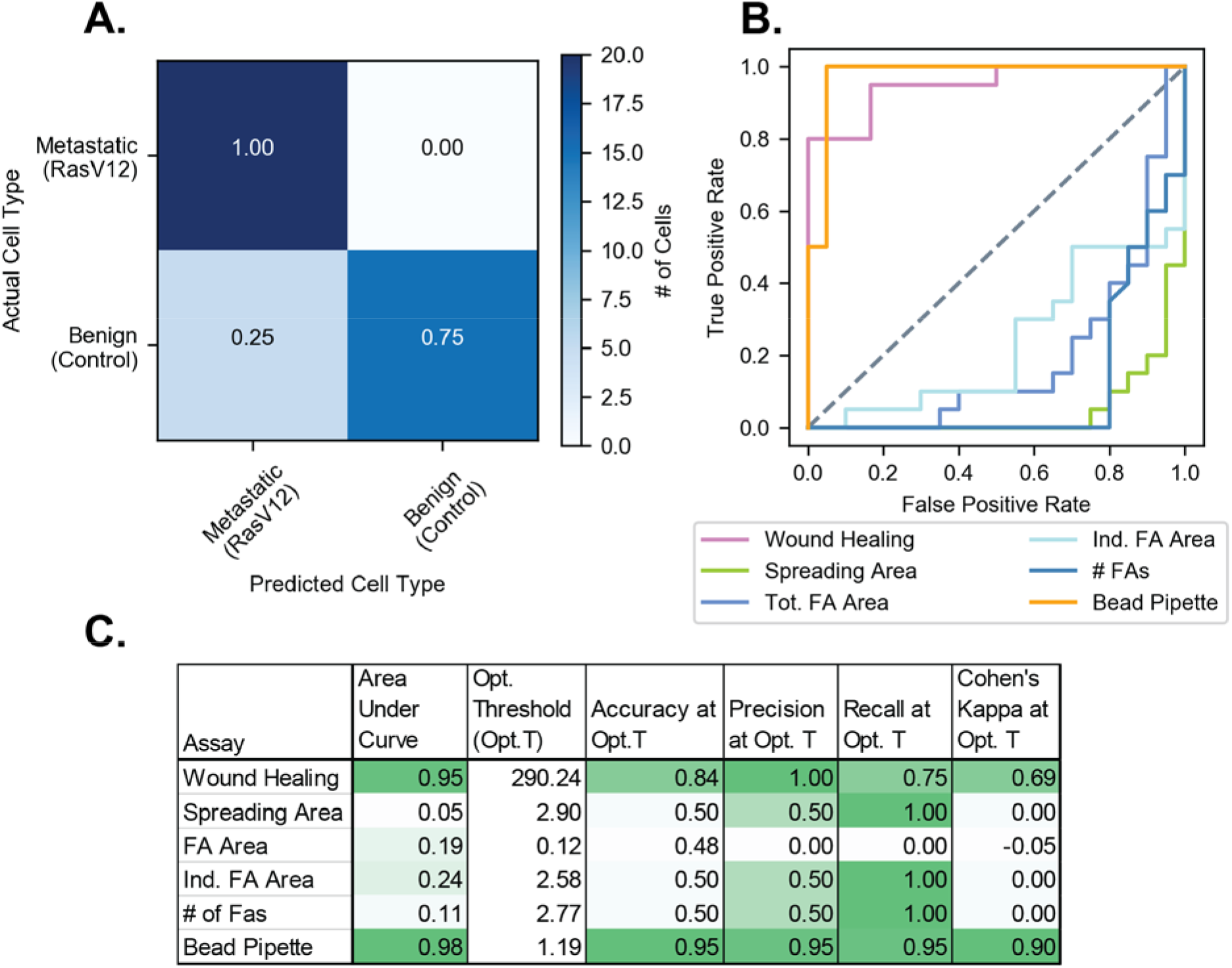
Statistical analysis of assays shows higher classifier performance of the bead-pipette assay. **A.** Example confusion matrix analysis for arbitrary bead pipette assay force threshold, plotted in Matplotlib(46). Cells were considered metastasizing if they exerted a force above the lower standard deviation value of the RasV12 cells (1.083 nN). Predictions were compared to cell genotypes. **B.** Receiver Operating Characteristic (ROC) curves for each assay, calculated with Scikit-Learn(45) and plotted in Matplotlib(46). **C.** Matrix comparing area under ROC curves and accuracy, precision, recall, and Cohen’s Kappa Coefficient(47) at the optimal threshold, determined through Youden’s J Statistic(48). Values were calculated with Scikit-Learn(45) and are colored relative to the range of values within each column.

To find the optimal force threshold and compare the bead pipette assay as a classifier against all other assays, ROC curves were plotted. The ROC curves in Fig 7B show that only the bead-pipette assay and the wound healing assay can classify the metastatic and benign cells at a standard above random classification. All other assays, while their differences are statistically significant in terms of the Student’s T-Test, perform lower than random classification. In Fig 7C, the area under the ROC curves show that the bead-pipette assay performs slightly higher than the wound healing, and both are markedly higher than other assays. Even at the optimal thresholds for each assay, the accuracy, precision, and Cohen’s Kappa - a statistic of classifier performance, while considering random classification(47) - are notably higher for the bead-pipette and wound healing assay. The only metric which the FA analyses and spreading area assays perform higher is the recall, however the other metrics indicate that although these assays can classify all the metastasizing cells correctly, they cannot effectively differentiate them from benign cells. The full confusion matrices for all assays at optimal threshold are in S1 Fig.

Overall, the bead-pipette assay performed well as a classifier compared to other assays that are often used to identify the hallmarks of metastatic cells. These statistical tests indicate that while the adhesion force model still requires more testing and refinement over larger sample sizes, using the bead-pipette assay to predict metastasis is promising.

## Conclusion

Assays in this work were able to identify several features of metastatic cells being expressed in the RasV12 NIH3T3 cells used to model metastatic cancer cells. The bead-pipette assay was able to quantify the difference between the control and RasV12 cells. The assay measured that the control cells exerted less force than RasV12 cells on the same substrate over the same contact time. The results also suggest that the turnover time may be significant in metastatic mechanisms, as over the same period of time the RasV12 cells, with smaller FAs, were able to adhere with more force, indicating a fast formation time.

The bead-pipette assay as a metastasis classifier has many advantages in this system. Concerning its potential as a clinical diagnostic, the assay uses widespread and inexpensive laboratory materials. To compare, AFM cantilevers and microscopes are expensive (on the order of tens of thousands of US dollars) and require specialized training, as well as can have technical drawbacks due to positional and time-based complications.

In addition, the bead-pipette assay measures individual cells, and only requires one seeding in order to get significant results. In other methods such as hydrodynamic shear or centrifugal force techniques, many cells are needed in order to produce significant results, as these techniques typically measure the force needed to displace 50% of the cells. However, since the bead-pipette assay measures each cell individually, fewer cells on the scale of a tumor biopsy would likely be sufficient.

With respect to research, the bead-pipette assay also has advantages for studying cell adhesion mechanisms. FA turnover time is an important factor in this work, however, most current techniques of force quantification detach cells with pre-established FAs. In contrast, the bead-pipette assay controls the contact time and is primarily invested in measuring nascent FAs; the assay gives insight into turnover rate.

In addition, the assay inflicts minimal deformation on the cell, allowing for repeated measurements over a period of time. A contrasting example is the micropipette aspiration technique, which displaces the cell from an adhered surface through suctioning the cell from a surface. However this method can tear the cell membrane and can only be performed once per cell. Other population-based methods also cannot repeat measurements on the same cells, as they displace a certain number of cells and may damage them while displacing them.

Lastly, the bead-pipette assay allows the researcher to observe the cell and perform the adhesion force measurement simultaneously. Although seemingly insignificant, this feature is not commonly available in all methods such as centrifugation, and may be important for observing particular phenotypes of interest. In this work, observing the cells from the measurement led to preliminary analysis, and also allowed recognition of cells in an undesirable growth phase, for example, apoptosis.

In conclusion, the novel bead-pipette assay has the potential to be a viable diagnostic tool for distinguishing patient metastatic cells based on adhesion force. The RasV12 cells have displayed multiple characteristics of metastasizing cells such as faster cell migration, polarized cell shape, smaller FA area, and less FA numbers compared to control cells. Unlike other methods, the bead-pipette assay is able to also account for turnover time of FA synthesis, and has shown that the RasV12 cells have faster turnover time to account for cell motility. Due to the simplicity of the technique and the novelty of the measurement, the bead-pipette assay has potential as an effective and accessible method of force quantification that applies a physical solution to a biological problem.

## Acknowledgements

This work was funded by the Mechanobiology Institute. In addition, the authors would like to acknowledge Professor Yan Jie for integral feedback and review on this work.

## Supporting Information

**S1 Fig. Confusion matrices for all assays at optimal thresholds.** Optimal thresholds and corresponding confusion matrices were calculated in Scikit-Learn(45) and plotted in Matplotlib(46).

**S1 Table. Wound healing assay data of cell migration over 8 hrs.**

**S2 Table. Spreading area assay data of cell area.**

**S3 Table. Focal Adhesion analyses data for Focal Adhesion area and size.**

**S4 Table. Bead-Pipette assay data for cell adhesion force.**

