## Supplementary figures and images for "Predicting metastasis with a novel biophysical cell-adhesion force technique"

### Supplemental Figure 1

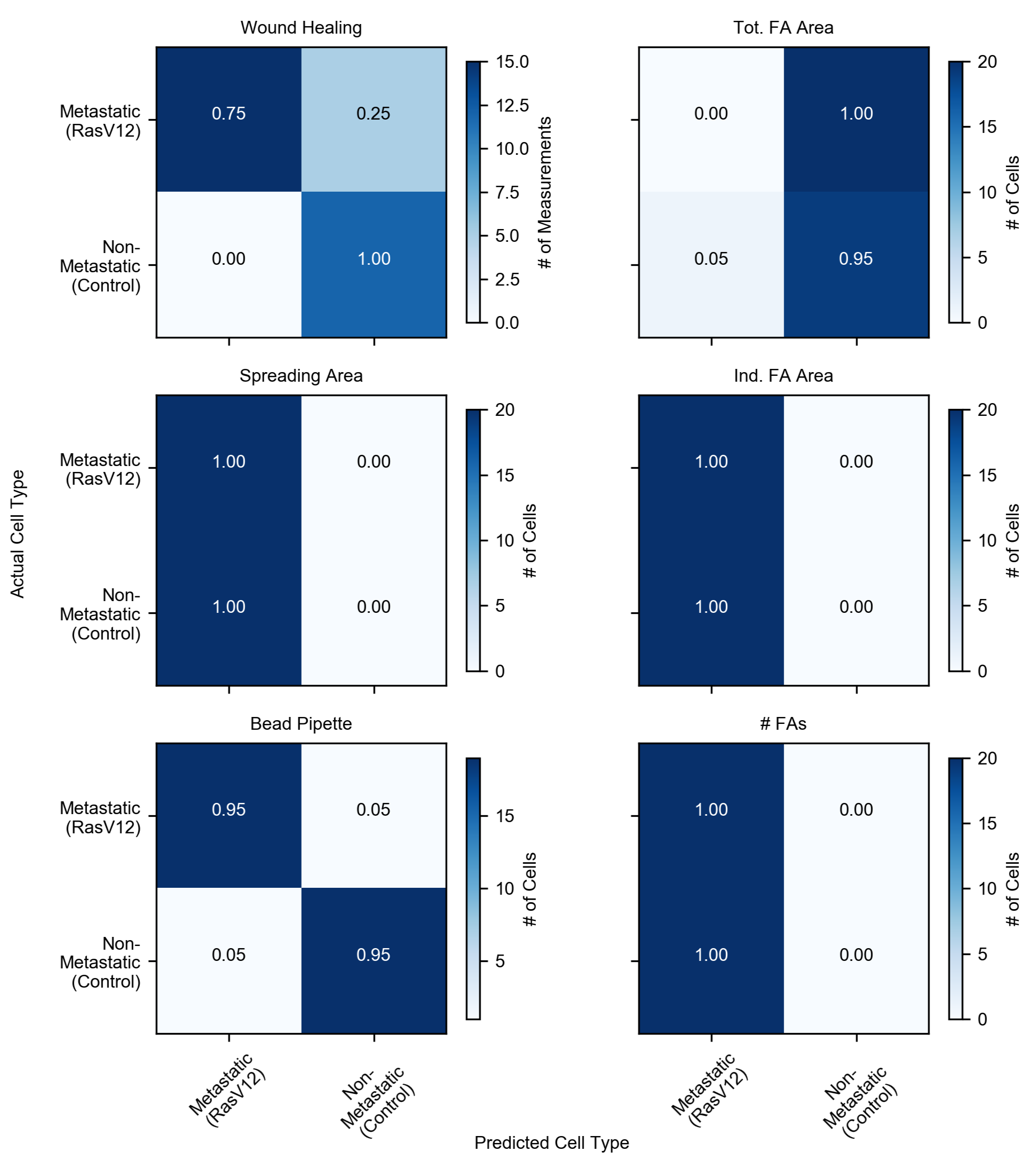
